# Thiazides induce glucose intolerance through inhibition of mitochondrial carbonic anhydrase 5b in β-cells

**DOI:** 10.1101/2022.03.23.485566

**Authors:** Patrycja Kucharczyk, Giuseppe Albano, Christine Deisl, Stephan Wueest, Daniel Konrad, Daniel G. Fuster

## Abstract

Thiazides are associated with glucose intolerance and new onset diabetes mellitus, but the molecular mechanisms remain elusive. The aim of this study was to decipher the molecular basis of thiazide-induced glucose intolerance. In mice, hydrochlorothiazide induced a pathological glucose tolerance, characterized by reduced first phase insulin secretion but normal insulin sensitivity. *In vitro*, thiazides inhibited glucose-and sulfonylurea-stimulated insulin secretion in islets and the murine β-cell line Min6 at pharmacologically relevant concentrations. Inhibition of insulin secretion by thiazides was CO_2_/HCO_3_^-^-dependent, not additive to unselective carbonic anhydrase (CA) inhibition with acetazolamide and independent of extracellular potassium. In contrast, insulin secretion was unaltered in islets of mice lacking the known molecular thiazide targets NCC (SLC12A3) or NDCBE (SLC4A8). CA expression profiling with subsequent knock-down of individual CA isoforms suggested mitochondrial CA5b as molecular target. In support of these findings, thiazides significantly attenuated Krebs cycle anaplerosis through reduction of mitochondrial oxalacetate synthesis. CA5b KO mice were resistant to thiazide-induced glucose intolerance, and insulin secretion of islets isolated from CA5b KO mice was unaffected by thiazides.

In summary, our study reveals attenuated insulin secretion due to inhibition of the mitochondrial CA5b isoform in β-cells as molecular mechanism of thiazide-induced glucose intolerance.

## INTRODUCTION

Arterial hypertension and kidney stones are highly prevalent and global health care problems (1, 2). Thiazide and thiazide-like diuretics (thiazides) have been the cornerstone for the treatment of arterial hypertension and pharmacologic recurrence prevention of kidney stones for more than 50 years. The preferred use of thiazides for these indications is supported by decades of clinical trials with hard clinical outcomes and real-world experience. Hence, not surprisingly, thiazides belong to the most widely prescribed drugs worldwide (3).

Since their introduction into clinical medicine in the 1960ies, thiazides are known to be associated with metabolic side effects, including glucose intolerance and new onset diabetes. In large prospective observational cohorts (4-6) and randomized controlled trials (7-9), thiazide use was associated with a significantly increased risk of new onset diabetes mellitus. Of all major classes of antihypertensive drugs, thiazides carry the highest risk for the development of diabetes (10). Several hypotheses have been put forth to explain thiazide-induced glucose intolerance, but the underlying mechanisms remain elusive until today (11). Unfortunately, these unpredictable and poorly understood side effects have caused many physicians to avoid the use of these clinically effective, ubiquitously available and cheap drugs. In recognition of this important knowledge gap in a clinically highly relevant area, a working group of the National Heart, Lung and Blood Institute issued a call for research on thiazide-induced dysglycemias a decade ago (11).

The classical molecular thiazide target is the Na^+^/Cl^-^ co-transporter NCC (also known as SLC12A3) in distal convoluted tubules (DCT) of the kidney. Inhibition of Na^+^ and Cl^-^ reabsorption by NCC is responsible for both the natriuretic and hypocalciuric action of the drug (12, 13). In addition to NCC, other molecular thiazide targets have been described. Thiazides were originally developed by chemical modification of carbonic anhydrase (CA) inhibitor acetazolamide (AZA) (14). Despite considerable chemical alterations of the original AZA scaffold, all thiazides have retained the ability to inhibit CA. However, there is considerable heterogeneity between the different thiazides with respect to their inhibitory potency towards the currently known CA isoforms (15, 16). Eladari and colleagues described an additional thiazide target, the Na^+^-driven Cl^-^/bicarbonate exchanger NDCBE (also known as SLC4A8) that mediates thiazide-sensitive Na^+^ transport in the cortical collecting duct (CCD) of the kidney (17).

The role of the currently known molecular thiazide targets in the development of thiazide-induced metabolic side effects remains unknown. The aim of this study was to decipher the molecular basis of thiazide-induced glucose intolerance.

## RESULTS

### Hydrochlorothiazide induces glucose intolerance in mice

In a first set of experiments, we assessed if thiazides induce glucose intolerance in mice. To this end, intraperitoneal glucose tolerance tests (IPGTT) were performed in 3 months old male C57BL/6J mice treated with intraperitoneally injected 1, 2, 5 or 50 mg/kg body weight hydrochlorothiazide (HCT) or vehicle. As shown in Fig. 1A-H, HCT treated mice exhibited significantly higher glycemic excursions during IPGTTs compared to vehicle treated mice in a dose-dependent manner. The lowest HCT dose associated with glucose intolerance was 5 mg/kg (Fig. 1 E,F). Blood glucose levels in 5 mg/kg and 50 mg/kg HCT treated mice were already higher compared to vehicle treated mice at time point 0 min, i.e. before the intraperitoneal glucose challenge but 30 minutes after HCT injection. After overnight fasting, IPGTT results were similar compared to 6 h fasted mice, but differences between vehicle and HCT treated mice only started to emerge after the intraperitoneal glucose challenge (Suppl. Fig 1). In all conditions, body weight was similar in HCT and vehicle treated mice (Fig. 1, Suppl. Fig. 1). In follow-up experiments, we measured serum insulin levels in mice treated with HCT or vehicle. As depicted in Fig. 1I-L, serum insulin levels were significantly lower 2 min after an intraperitoneal glucose challenge (2 g/kg body weight) in mice pre-treated with either 5 or 50 mg/kg body weight HCT, indicating reduced first-phase insulin secretion in HCT treated mice. To assess the impact of HCT on insulin sensitivity, we performed intraperitoneal insulin tolerance tests (IPITT). As demonstrated in Fig. 1M-P, insulin sensitivity, tested by two different insulin doses (0.5 IU/kg and 1 IU/kg body weight, respectively), was not altered by 5 mg/kg body weight HCT application. Together these findings suggest reduced insulin secretion but maintained insulin sensitivity as potential mechanism for HCT-induced glucose intolerance. In addition, the results obtained support the use of mice as a mammalian model organism to further investigate the molecular mechanisms of thiazide-induced on glucose intolerance.

**Figure 1.**
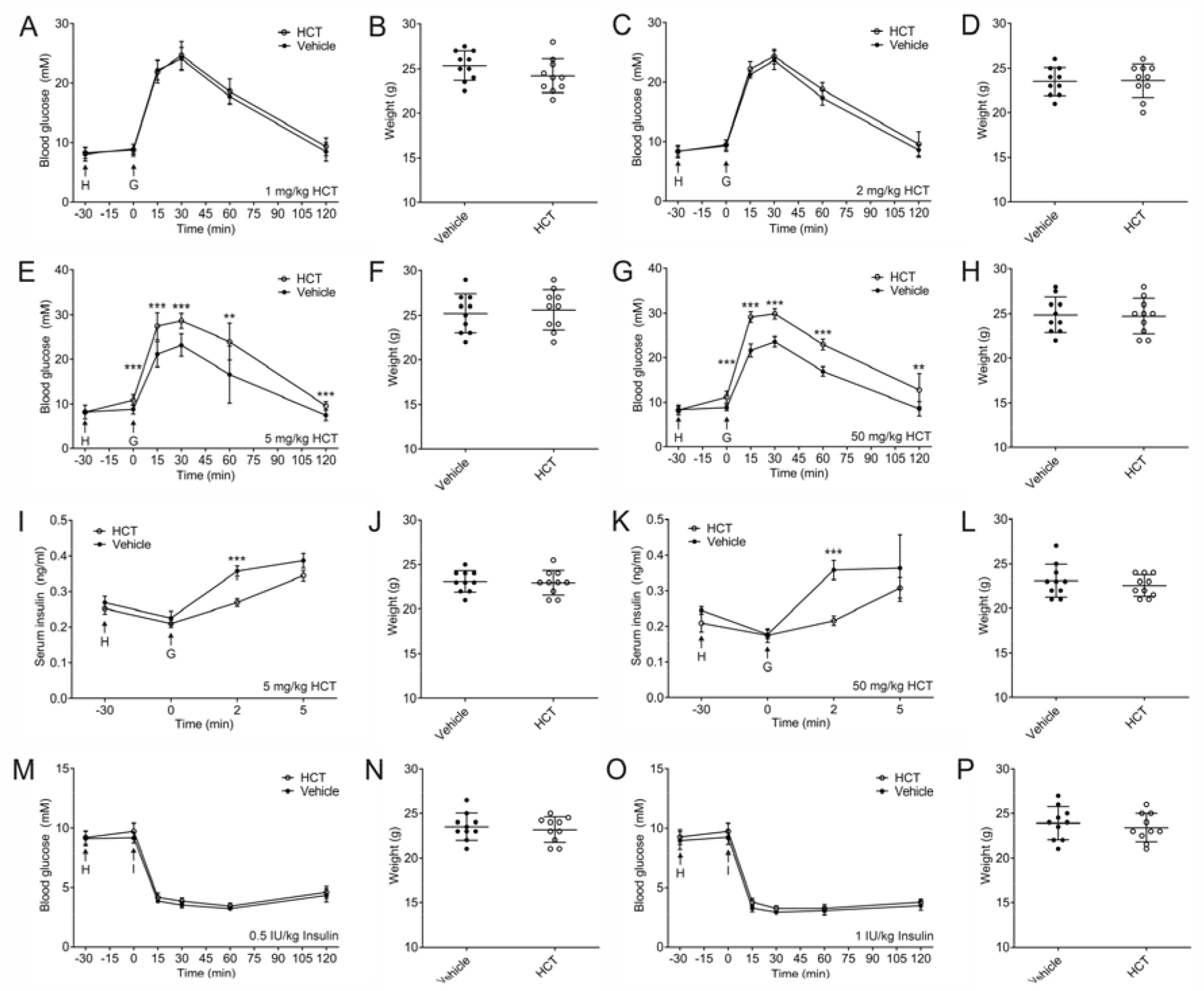
Hydrochlorothiazide induces glucose intolerance in mice. Intraperitoneal glucose tolerance tests and corresponding body weights of C57BL/6J mice treated with 1 mg/kg **(A** and **B)**, 2 mg/kg **(C** and **D)**, 5 mg/kg **(E** and **F)** and 50 mg/kg **(G** and **H)** hydrochlorothiazide (HCT) or vehicle in 6 h fasted mice. HCT or vehicle i.p. (arrow H) was applied at time point -30 min, glucose (2g/kg; arrow G) was applied at time point 0 min. Whole blood glucose was measured at indicated time points. Serum insulin and corresponding body weights of C57BL/6J mice treated with 5 mg/kg **(I** and **J)** or 50 mg/kg **(K** and **L)** HCT or vehicle i.p. (arrow H) at time point -30 min and glucose (2g/kg; arrow G) at time point 0 min. Intraperitoneal insulin tolerance tests and corresponding body weights of C57BL/6J mice treated with 5 mg/kg HCT or vehicle i.p. (arrow H) at time point -30 min and insulin (arrow I) at time point 0 min. **(M** and **N)** insulin 0.5 IU/kg i.p., **(O** and **P)** insulin 1 IU/kg i.p. N=10 mice per group in all experiments. Data are show as mean ± SD. Asterisks denote significance for comparisons between groups of mice at indicated time points (two-tailed unpaired Student’s t-test; **p <0.01, ***p <0.001).

In humans, robust diuretic effects are observed with HCT doses of 1-2 mg/kg body weight (18, 19). Significantly higher thiazide doses are needed in mice to stimulate natriuresis, in case of HCT typically doses of 20 – 50 mg/kg body weight are applied (20-28). The reason for this discrepancy is not entirely clear, but likely due to differences in the pharmacokinetics of thiazides between mice and humans. To this end, we established the dose-response for the natriuretic effect of HCT in mice, which reflects inhibition of the primary thiazide target NCC in the DCT. As shown in Suppl. Fig. 2, there was a dose-dependent increase in urinary Na^+^ excretion. The 5 mg/kg but not 1 mg/kg HCT dose increased urinary Na^+^ excretion compared to vehicle treated mice, and the natriuretic effect was higher with 50 mg/kg compared to 5 mg/kg. Hence, the 5 mg/kg HCT dose in mice corresponds to a HCT dose typically employed in humans (12.5 – 25 mg daily) for the treatment of arterial hypertension (29). In case of recurrence prevention of calcium kidney stones, even higher doses (50 – 100 mg daily) are employed (30).

### Thiazides attenuate insulin secretion in islets and Min6 cells

We then assessed the impact of HCT on insulin secretion in primary islets isolated from C57BL/6J mice and in the murine β-cell line Min6 (Fig. 2A,B, Suppl. Fig. 3). HCT significantly attenuated insulin secretion in islets and Min6 cells *in vitro* at pharmacologically relevant concentrations (10^−7^ – 10^−9^ M), while not affecting cell viability (31). We obtained similar findings with other thiazides, including metolazone, chlorthalidone, indapamide and bendroflumethiazide (Suppl. Fig. 4). HCT also significantly lowered sulfonylurea-induced insulin secretion (250 μM tolbutamide) while it only had a small effect on basal (2 mM glucose) and direct depolarization-induced insulin secretion by high (30 mM) extracellular K^+^ (Fig. 2C) (32, 33).

**Figure 2.**
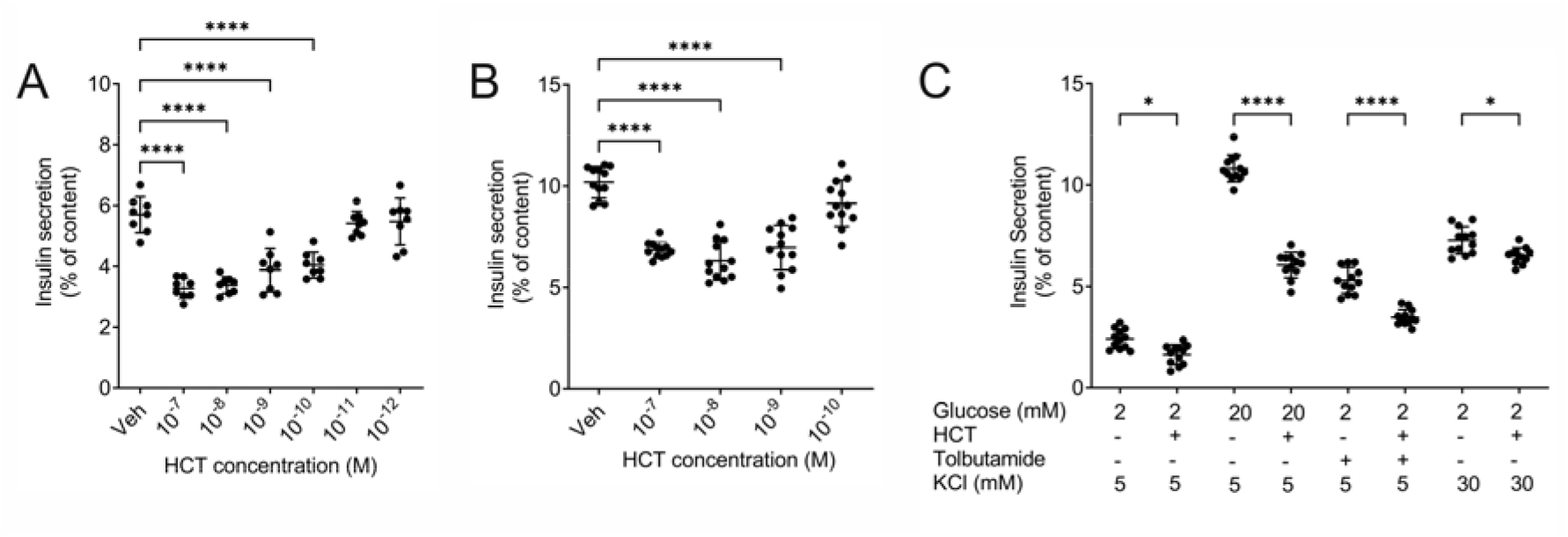
Hydrochlorothiazide inhibits insulin secretion in murine islets and Min6 cells. **(A)** Glucose-stimulated insulin secretion (20 mM glucose; GSIS) of islets isolated from C57BL/6J mice incubated with vehicle (DMSO 1:1000) or hydrochlorothiazide (HCT) at indicated concentrations. Each dot represents islets isolated of an individual mouse. **(B)** GSIS of Min6 cells incubated with vehicle or HCT at indicated concentrations. Data represent three individual experiments combined. **(C)** Insulin secretion of Min6 cells in the presence of indicated concentrations of glucose (2 or 20 mM) and KCl (5 or 30 mM), treated with 10^−8^ M HCT or 250 μM tolbutamide. Data represent three individual experiments combined. Data are shown as individual observations with mean ± SD. Asterisks denote significance for the indicated comparisons (ANOVA with Tukey post-hoc test; *p <0.05, ****p < 0.0001).

### Thiazides target carbonic anhydrase in β-cells

To further define the basis of reduced insulin secretion by thiazides, we assessed mRNA expression of the known thiazide targets including NCC, NDCBE and CA isoforms in Min6 cells and murine islets (Suppl. Fig. 5A-D). Transcripts of both NDCBE and NCC were detectable in Min6 cells and islets. Of the 15 CA isoforms CA1, CA4 and CA5a were not detectable in both Min6 cells and islets. CA6 and CA7 were expressed in Min6 cells but not islets. We next assed the impact of genetic NDCBE or NCC deletion in islets on insulin secretion. As shown in Suppl. Fig. 5E,F, islets isolated of NDCBE and NCC knock-out (KO) mice displayed no insulin secretion deficit.

In humans, biallelic pathogenic variants in *SLC12A3*, encoding NCC, result in Gitelman’s syndrome which is characterized by hypotension, hypokalemia, hypomagnesemia and metabolic alkalosis (34, 35). In addition to the well-known electrolyte abnormalities, patients affected by Gitelman’s syndrome were reported to exhibit an increased prevalence of impaired glucose tolerance (36-38). To assess the role of NCC in systemic glucose homeostasis, we additionally performed IPGTTs in NCC KO mice. As shown in Supp. Fig. 5G, H, genetic loss of NCC was not associated with altered glucose tolerance.

We then assessed the role of CA in insulin secretion. Inhibition of CA with the non-specific CA inhibitor AZA significantly attenuated insulin secretion in both Min6 cells and islets *in vitro* (Fig. 3A,B) and induced glucose intolerance *in vivo* (Fig. 3C,D). Compared to either AZA or HCT alone, the combination of AZA and HCT did not further attenuate insulin secretion (Fig. 3 E), and HCT had no effect on insulin secretion in a CO_2_/HCO_3_ ^-^ free condition (Fig. 3 F). These results reveal that the effect of HCT on insulin secretion is CO_2_/HCO_3_^-^ -dependent and not additive to (unselective) CA inhibition, suggesting that HCT targets one or several CA isoform(s) in β-cells that are critical for insulin secretion.

**Figure 3.**
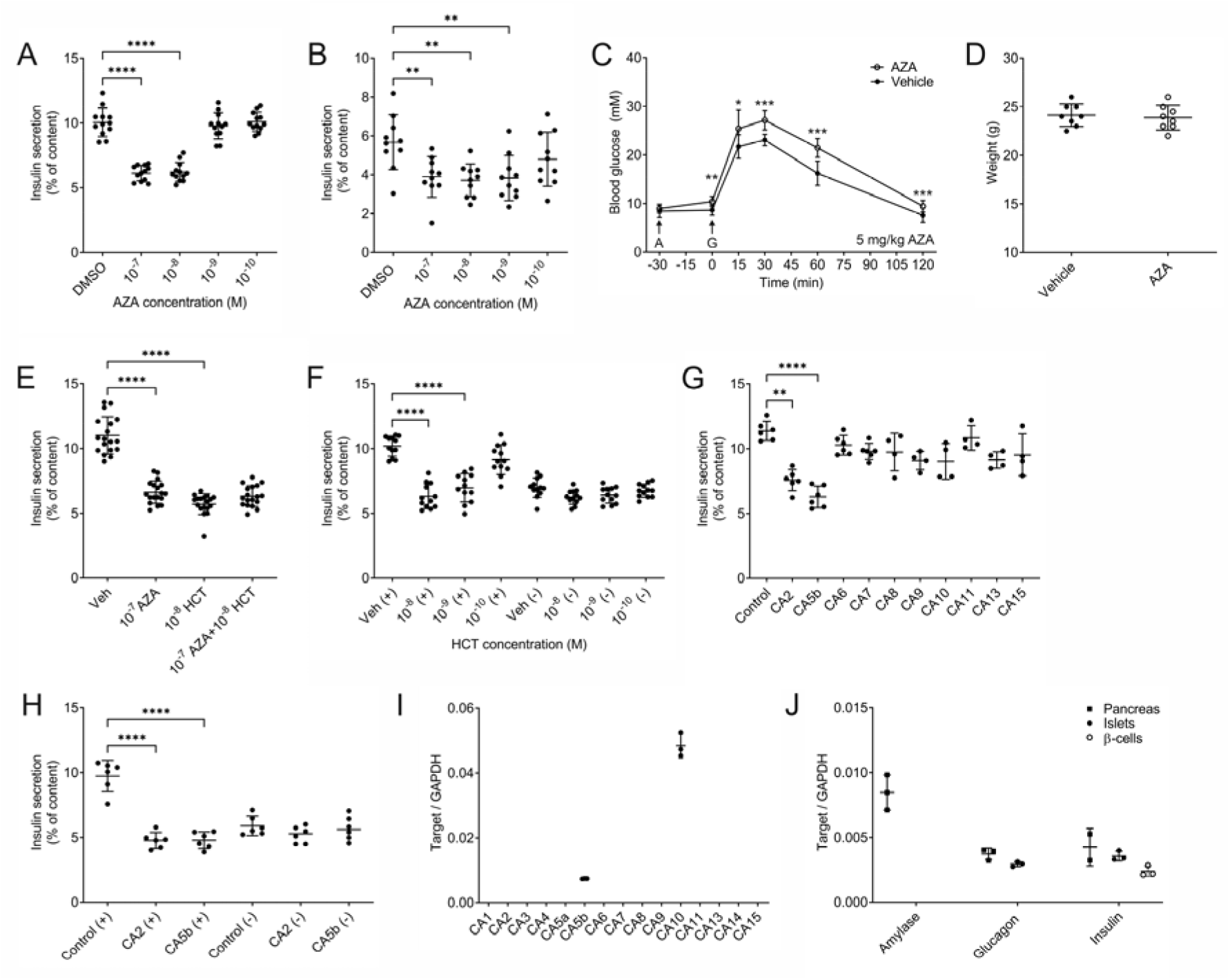
Hydrochlorothiazide targets carbonic anhydrase in β-ells. **(A)** Glucose-stimulated insulin secretion (20 mM glucose; GSIS) of Min6 cells incubated with vehicle (DMSO 1:1000) or the carbonic anhydrase inhibitor acetazolamide (AZA) at indicated concentrations. **(B)** GSIS of islets isolated from C57BL/6J mice incubated with vehicle or AZA at indicated concentrations. Each dot represents islets isolated of an individual mouse. **(C** and **D)** Intraperitoneal glucose tolerance tests and corresponding body weights of C57BL/6J mice treated with 5 mg/kg AZA. AZA or vehicle i.p. (arrow A) was applied at time point – 30 min, glucose (2g/kg; arrow G) was applied at time point 0 min. Whole blood glucose was measured at indicated time points. N=8 mice per group. **(E)** GSIS of Min6 cells treated with vehicle or indicated concentrations (M) of either AZA or HCT, or AZA and HCT combined. **(F)** GSIS of Min6 cells in the presence (+) or absence (-) of CO_2_/HCO_3_ ^-^, treated with vehicle or HCT at indicated concentrations. **(G)** GSIS of Min6 cells treated with control siRNA or siRNAs targeting indicated carbonic anhydrase (CA) isoforms. **(H)** GSIS of Min6 cells treated with control siRNA or siRNAs targeting CA2 or CA5b, respectively, in the presence (+) or absence (-) of CO_2_/HCO_3_ ^-^. **(I)** Expression profiling of CA isoforms in purified murine β-cells by real-time PCR and normalized to GAPDH. Each dot represents β-cells isolated from an individual mouse. **(J)** Assessment of purity of isolated islets and purified β-cells by transcript expression analysis of amylase (marker of exocrine pancreas), glucagon (marker of α-cells) and insulin (marker of β-cells) in pancreas, islets and purified β-cells by real-time PCR and normalized to GAPDH. Each dot represents tissue or cells isolated from an individual mouse. Data are shown as individual observations with mean ± SD. Min6 cell experiments represent three individual experiments combined. Asterisks denote significance for the indicated comparisons of *in vitro* insulin secretion studies (ANOVA with Tukey post-hoc test; **p <0.01, ****p < 0.0001) or for comparisons between two groups of mice at indicated time points (two-tailed unpaired Student’s t-test; **p < 0.05, **p < 0.01, ***p < 0.01).

Currently, there are no specific CA isoform inhibitors available. To identify the responsible CA isoform(s), we performed siRNA-mediated knock-down of all CA isoforms expressed in Min6 cells (Fig. 3G, Suppl. Fig. 5C, Suppl. Fig. 6) and performed static insulin secretion experiments. Only CA2 and CA5b knock-down significantly attenuated insulin secretion in Min6 cells, whereas the other CA isoforms expressed in Min6 cells were dispensable for insulin secretion (Fig. 3G). In addition, omission of CO_2_/HCO_3_ ^-^ in the incubation medium did not further impair insulin secretion in either CA2 or CA5b depleted cells compared to control, indicating critical but non-redundant roles of these two CA isoforms for insulin secretion in Min6 cells (Fig. 3H). We then assessed CA isoform mRNA expression in murine islets and purified β-cells. As shown in Fig. 3I, J and Suppl. Fig. 5D, while many CA isoforms are expressed in islets, β-cells only express the two isoforms CA5b and CA10. These findings, together with the fact that CA10 is a catalytically inactive and secreted CA isoform, suggest mitochondrial CA5b as the likely target of thiazides in β-cells (39).

### Hydrochlorothiazide inhibits Krebs cycle anaplerosis in β-cells

Mitochondria are impermeant to HCO_3_^-^, yet several mitochondrial enzymes critically depend on HCO_3_^-^ such as pyruvate carboxylase, which generates oxaloacetate (OA) from pyruvate and HCO_3_ ^-^ (40). The two mitochondrial CA isoforms CA5a and CA5b utilize CO_2_, which freely diffuses into mitochondria, to produce HCO_3_^-^ needed for these enzymatic reactions. β-cells exhibit high pyruvate carboxylase (PC) activity but do not perform gluconeogenesis because they lack phosphoenolpyruvate carboxykinase. However, PC restores Krebs cycle intermediates lost during metabolic reactions by producing OA from pyruvate (anaplerosis) (41-43).

In a next step, we measured glucose-induced insulin secretion in Min6 cells in the presence of HCT, the PC inhibitor phenylacetic acid (PAA) or a combination of both, with and without CO_2_/HCO_3_^-^. As shown in Fig. 4A, inhibition of glucose-stimulated insulin secretion was equal with incubation of HCT or PAA, and the combination of both did not result in a further reduction. Furthermore HCT, PAA or the combination of both had no effect on glucose-stimulated insulin secretion in CO_2_/HCO_3_ ^-^-free conditions. Similarly, in murine islets, glucose-stimulated insulin secretion was reduced to a similar degree in the presence of HCT or PAA, and the combination of both HCT and PAA did not result in a further reduction (Fig. 4B).

**Figure 4.**
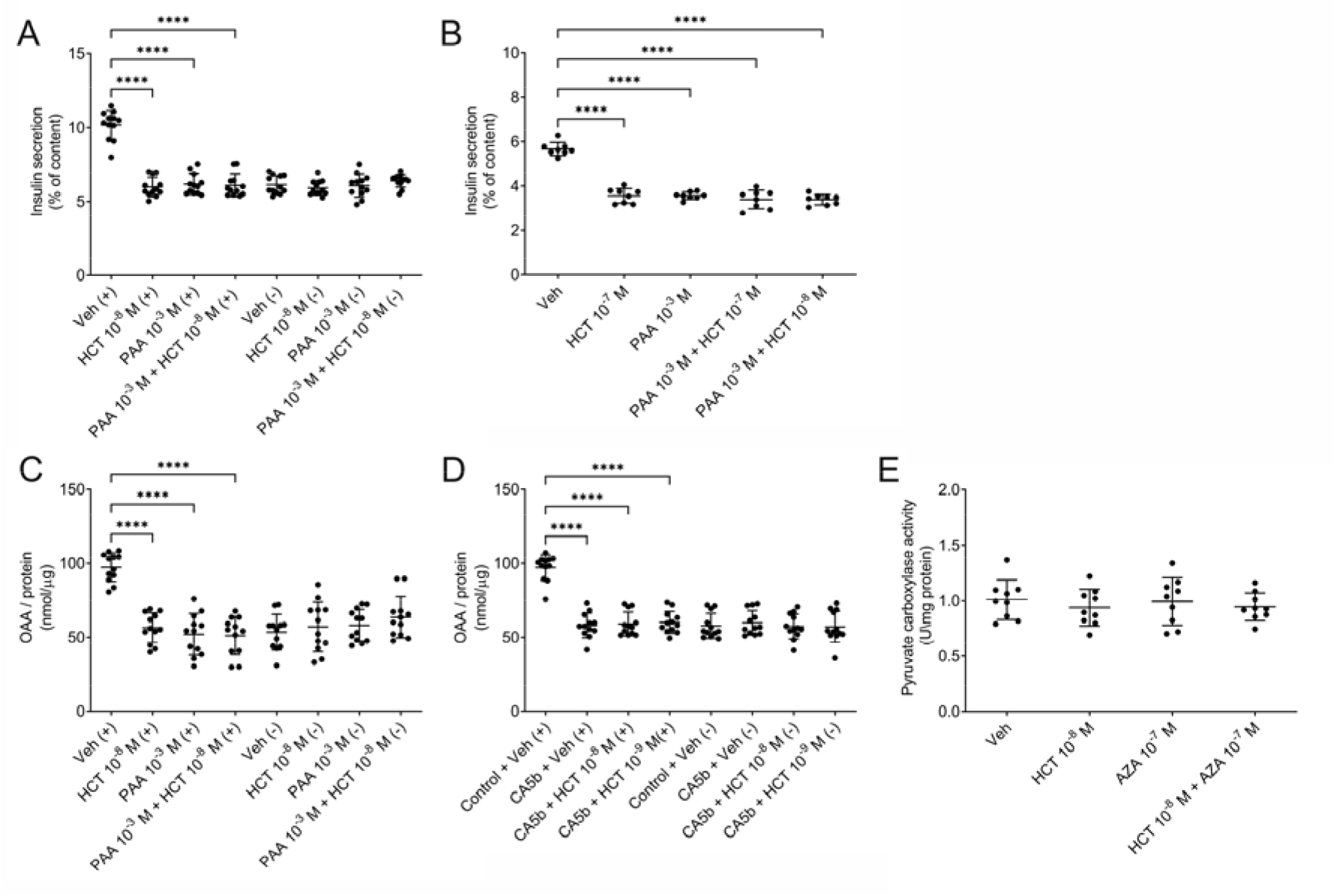
Hydrochlorothiazide attenuates oxalacetate synthesis in β-cells. **(A)** Glucose-stimulated insulin secretion (20 mM glucose; GSIS) of Min6 cells in the presence (+) or absence (-) of CO_2_/HCO_3_^-^, treated with vehicle (DMSO 1:1000), hydrochlorothiazide (HCT), the pyruvate carboxylase inhibitor phenylacetic acid (PAA) or a combination of PAA and HCT at indicated concentrations. **(B)** GSIS of islets isolated from C57BL/6J mice incubated vehicle, HCT, PAA or a combination of PAA and HCT at indicated concentrations. **(C)** Oxalacetate (OAA) content normalized to protein of Min6 cells stimulated with 20 mM glucose, in the presence (+) or absence (-) of CO_2_/HCO_3_ ^-^ and treated with either vehicle, HCT, PAA or a combination of HCT and PAA at indicated concentrations. **(D)** OAA content normalized to protein of Min6 cells stimulated with 20 mM glucose, in the presence (+) or absence (-) of CO_2_/HCO_3_^-^, treated with control siRNA or siRNA targeting CA5b, and incubated with either vehicle or HCT at indicated concentrations. **(E)** Pyruvate carboxylase activity in lysates of Min6 cells cultured for 4 h in the absence of CO_2_ prior to lysis, measured in CO_2_-free condition in the presence of indicated concentrations of HCT or AZA. Data are shown as individual observations with mean ± SD and represent three individual experiments combined. Asterisks denote significance for the indicated comparisons (ANOVA with Tukey post-hoc test; ****p < 0.0001).

We then measured OA levels in Min6 cells exposed to HCT or PAA during glucose stimulation. As shown in Fig. 4C, both HCT and PAA attenuated OA levels in Min6 cells by a similar magnitude and the combination of both did not result in a further reduction of OA levels. Furthermore HCT, PAA or the combination of both had no effect on OA levels in Min6 cells in CO_2_/HCO_3_^-^-free conditions. Finally, we treated Min6 cells with control or CA5b siRNA, and measured OA in the presence and absence of CO_2_/HCO_3_^-^. CA5b depletion or HCT were equally effective in reducing OA levels in Min6 cells, and this effect was again clearly CO_2_/HCO_3_^-^-dependent (Fig. 4D).

Due to the tight functional coupling between CA5b and PC, a direct effect of thiazides on PC activity cannot be definitively ruled out with studies employing intact cells. To test for a possible effect of HCT on PC, we performed *in vitro* PC activity experiments in Min6 cell lysates in the absence of CO_2_ with exogenous administration of HCO_3_^-^. As demonstrated in Fig. 4E, HCT, AZA or the combination of HCT and AZA had no effect on PC activity, further supporting the notion of CA5b as molecular target of HCT in β-cells. To substantiate this claim, we performed IPGTTs in CA5b KO and WT littermate mice (mixed C57BL6/J / SV129 background) treated with 5mg/kg body weight HCT or vehicle (40). HCT-treated WT mice exhibited significantly higher glycemic excursions during IPGTTs compared to vehicle treated mice (Fig. 5A,B). In contrast, we observed no difference between HCT and vehicle treated CA5b KO mice (Fig. 5C,D). Furthermore, in WT but not CA5b KO mice, HCT treatment was associated with lower serum insulin after the glucose challenge compared to vehicle treatment (Fig. 5E-H). We then performed IPITTs to assess the impact of HCT on insulin sensitivity in 3 months old male WT and CA5b KO mice. As demonstrated in Figs 5I,K, insulin sensitivity was similar in all groups of mice. Together, these *in vivo* findings demonstrate that genetic deletion of CA5b confers resistance to HCT-induced glucose intolerance.

**Figure 5.**
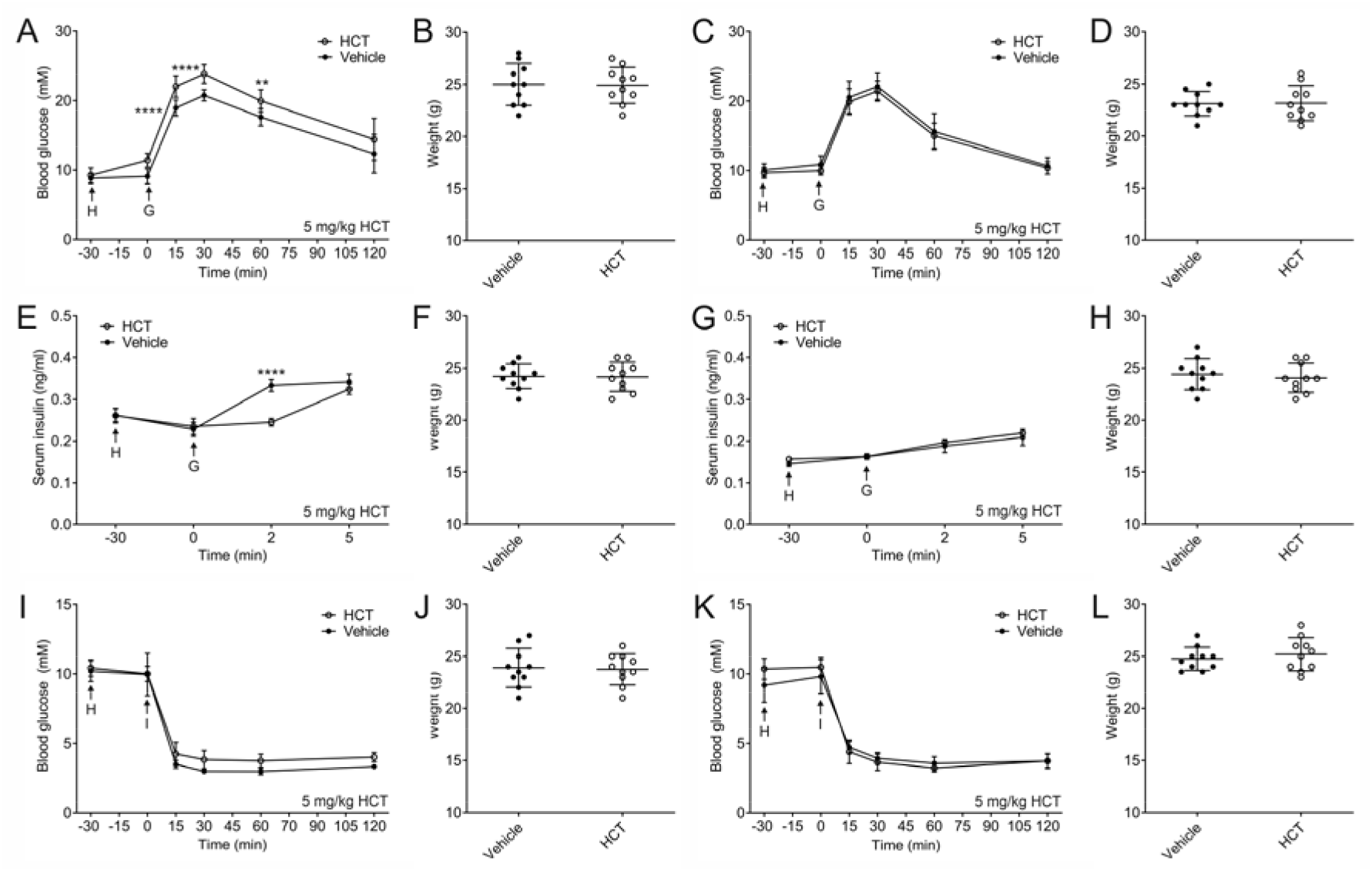
CA5b KO mice are resistant to hydrochlorothiazide-induced glucose intolerance. Intraperitoneal glucose tolerance tests in WT **(A** and **B)** and CA5b KO **(C** and **D)** mice with corresponding body weights treated with 5 mg/kg hydrochlorothiazide (HCT). HCT or vehicle i.p. (arrow H) was applied at time point – 30 min, glucose (2g/kg; arrow G) was applied at time point 0 min. Whole blood glucose was measured at indicated time points. Serum insulin in WT **(E** and **F)** and CA5b KO **(G** and **H)** mice with corresponding body weights treated with 5 mg/kg HCT or vehicle i.p. (arrow H) at time point – 30 min and insulin (2g/kg; arrow G) at time point 0 min. Intraperitoneal insulin tolerance tests in WT **(I** and **J)** and CA5b KO **(K** and **L)** mice with corresponding body weights treated with 5 mg/kg HCT or vehicle i.p. (arrow H) at time point – 30 min and insulin (1 IU/kg i.p.; arrow I) at time point 0 min. N=10 mice per group in all experiments. Data are show as mean ± SD. Asterisks denote significance for the indicated comparisons (two-tailed unpaired Student’s t-test; ****p < 0.0001).

To assess the role of genetic CA5b deletion on insulin secretion, we performed static insulin secretion experiments with islets isolated from WT and CA5b KO mice (Fig. 6A, Suppl. Fig. 7A, B). No CA5b expression was detectable in islets isolated from CA5b KO mice (Suppl. Fig. 7A). Furthermore, expression profiling of other CA isoforms did not reveal differences between WT and CA5b KO islets (Suppl. Fig. 7B). As previously observed with islets isolated from C57BL/6J mice, HCT attenuated insulin secretion in islets isolated from WT littermates of CA5b KO mice (mixed C57BL/6J/SV129 background). While insulin secretion of CA5b KO islets was significantly reduced compared to WT islets in the presence of the vehicle, HCT had no effect on insulin secretion of CA5b KO islets. A similar pattern was observed when we quantified OAA levels in WT and CA5b KO islets (Fig. 6B). In WT islets, HCT attenuated both high glucose (20 mM)- and sulfonylurea (250 μM tolbutamide)-induced insulin secretion, but did not influence basal (2mM glucose) or direct depolarization-induced insulin secretion by high (30 mM) extracellular K^+^ (Fig. 6C). In contrast, HCT did not affect basal or stimulated insulin secretion in CA5b KO islets (Fig. 6D). Together these results demonstrate that mitochondrial OA synthesis and insulin secretion capacity of CA5b KO islets are resistant to the action of HCT.

**Figure 6.**
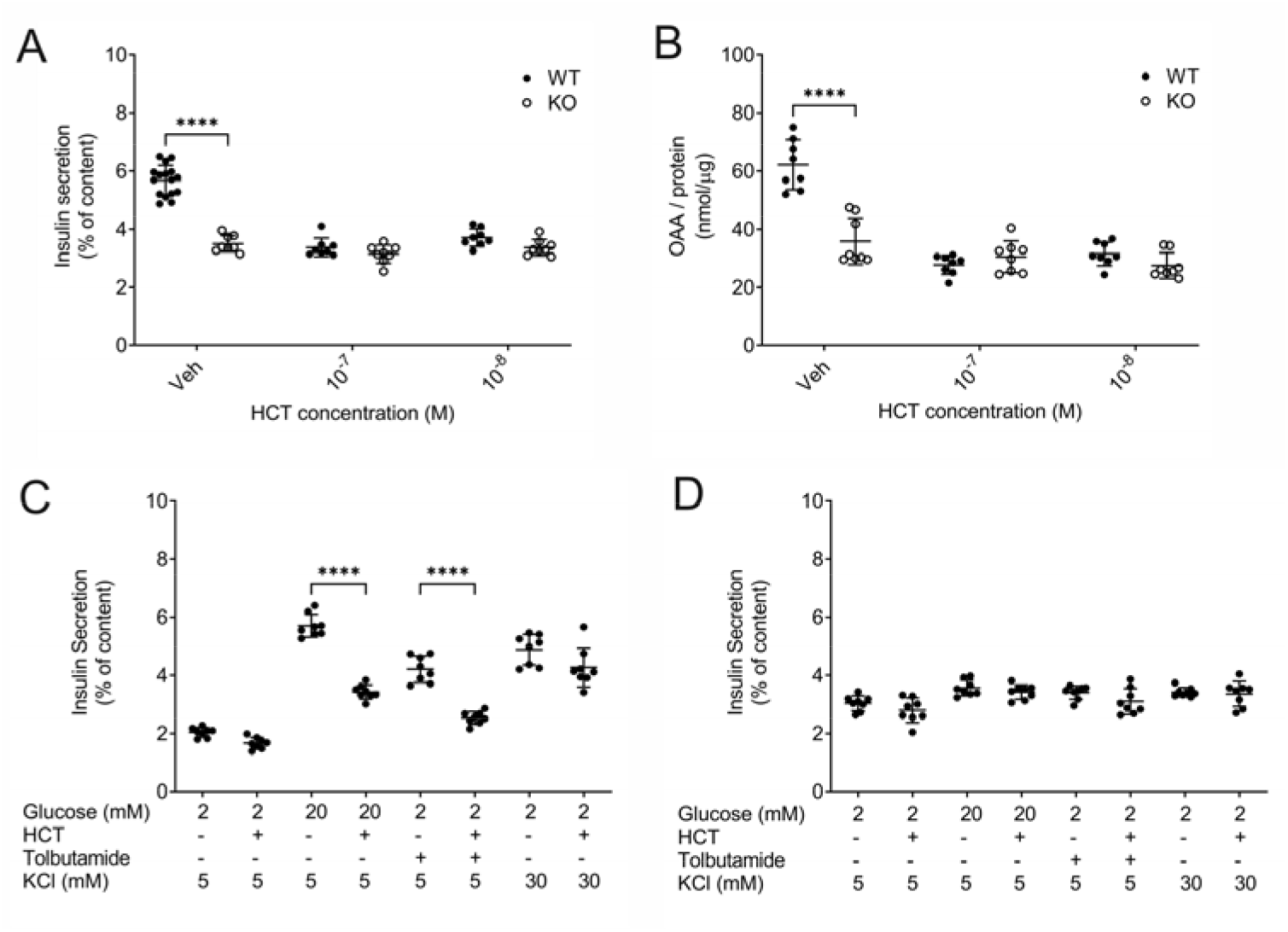
CA5b KO islets do not respond to classical secretagogues and are resistant to hydrochlorothiazide. **(A)** Glucose-stimulated insulin secretion (20 mM glucose) of WT (filled circles) and CA5b KO (open circles) islets treated with vehicle (DMSO 1:1000) or hydrochlorothiazide (HCT) at indicated concentrations. **(B)** Oxalacetate content normalized to protein of WT (filled circles) and CA5b KO (open circles) islets stimulated with 20 mM glucose and incubated with vehicle or HCT at indicated concentrations. Insulin secretion of WT **(C)** or CA5b KO **(D)** islets in the presence of indicated concentrations of glucose (2 or 20 mM) and KCl (5 or 30 mM), incubated with vehicle, HCT (10^−8^ M) or tolbutamide (250 μM). Data are shown as individual observations with mean ± SD. Each dot represents islets isolated from an individual mouse. Asterisks denote significance for the indicated comparisons (ANOVA with Tukey post-hoc test; ****p < 0.0001).

## DISCUSSION

Our study reveals that thiazides induce acute glucose intolerance in mice via attenuation of insulin secretion through inhibition of CA5b in β-cells. HCT and other frequently used thiazides, such as chlorthalidone, indapamide, metolazone and bendroflumethiazide, inhibited insulin secretion in a pharmacologically relevant, submicromolar range. CA expression profiling in Min6 cells and islets and subsequent siRNA knock-down experiments of individual CA isoforms in Min6 cells suggested the mitochondrial CA5b isoform as molecular target of thiazides in β-cells. In support of these results, CA5b KO mice were resistant to HCT-induced glucose intolerance, and insulin secretion of CA5b-deficient islets or Min6 cells was unaffected by HCT.

Mitochondrial CA5b provides HCO_3_^-^ for anaplerotic OA synthesis from pyruvate by PC. Deletion of CA5b or treatment with HCT attenuated OA synthesis in islets or Min6 cells. In line with these findings, purified full length (human) CA5b was previously shown to be inhibited by HCT and other thiazides *in vitro* with a K_i_ in the submicromolar range (15). While direct PC inhibition by PAA in islets mimicked the findings obtained with HCT treatment or CA5b deletion, our *in vitro* studies in CO_2_-free conditions with exogenous administration of HCO_3_^-^ demonstrate that PC is not inhibited by HCT or other thiazides up to the concentration of 10^−5^ M. This experiment enabled to functionally separate the two closely interacting enzymes CA5b and PC and led to the conclusion that PC *per se* is not thiazide-sensitive.

Cataplerotic export of Krebs cycle intermediates to the cytosol is critical for the amplification of glucose-and sulfonylurea-induced insulin secretion (41-43). In support of this, we found that secretagogue-induced insulin secretion was severely impaired in CA5b KO islets. Attenuation of insulin secretion was much more pronounced in CA5b KO islets compared to acute treatment with thiazides or knock-down with siRNA. While basal insulin secretion in low glucose condition was increased in CA5b KO islets compared to WT islets, CA5b KO islets were completely unresponsive to stimulation by high glucose, tolbutamide and high extracellular K^+^. In contrast, acute inhibition of PC, treatment with thiazides or knock-down of CA5b attenuated but not completely abrogated secretagogue-induced insulin secretion in Min6 cells and islets. In Min6 cells but not WT islets, HCT also slightly reduced basal and high K^+^-stimulated insulin secretion. The reason for these differences is not clear at the moment. We speculate that residual CA5b activity (inhibition, siRNA knock-down) versus complete loss of CA5b activity (KO model), and acute (inhibition, siRNA knock-down) versus chronic CA5b deficiency (KO model) may play a role. Certainly, a limitation of our study is the lack of a conditional CA5b KO mouse model that could be used to further explore this question.

Interestingly, Min6 cells express several CA isoforms, and seem to depend on cytosolic CA2 activity in addition to mitochondrial CA5b for insulin secretion. In contrast, primary murine β-cells only express CA5b and CA10. While CA2-positive pancreatic cells are progenitors of both exocrine and endocrine pancreatic cells, CA2 remains highly expressed in the exocrine but not endocrine pancreas upon differentiation (44). Likely, immortalization procedures, clonal selection artifacts and/or adaptations to cell culture conditions are responsible for the altered CA expression profile and the acquired dependence on CA2 for insulin secretion in Min6 cells. As is the case for CA5b, also CA10 is expressed in both Min6 cells and primary β-cells. CA10 is a catalytically inactive, secreted glycoprotein that was recently shown to physically interact with neurexins, a family of presynaptic adhesion molecules, and to facilitate their surface transport (39, 45). Hence, our data intriguingly suggest that β-cells do not express a cytoplasmic CA isoform. Furthermore, only one of the two mitochondrial CA isoforms is expressed in β-cells. CA5b KO mice do not exhibit an overt phenotype, normal growth and no evidence of metabolic abnormalities (40). In contrast, CA5a knock-out mice exhibit reduced growth, poor fertility and hyperammonemia as a result of defective ureagenesis in the liver. This suggests non-redundant physiological roles of the two mitochondrial CA isoforms. Purified CA5a can also be inhibited by thiazides *in vitro*, but the K_i_ values for thiazides are much higher compared to CA5b (15). Nevertheless, an effect of thiazides on CA5a activity with therapeutic doses of thiazides would theoretically still be possible and should be investigated further.

HCT induced hyperglycemia in mice already before the administration of a glucose challenge, but without discernable differences in serum insulin. Interestingly, this phenomenon was absent in mice fasted overnight or in mice lacking CA5b. We have no definitive explanation for the observation but speculate that the hyperglycemia is indeed mediated by a reduction in insulin secretion in the 30 min after HCT administration (which we missed) and that there is already a compensatory reduction in hepatic insulin clearance at time point 0. Other possible scenarios include a temporary increase in glucose elevating hormones (such as glucagon) or a temporary increase in hepatic glucose output by HCT. While the definitive mechanism remains to be established, our data clearly demonstrate that the increase in blood glucose observed prior to the glucose challenge by HCT is also dependent on CA5b.

In the past, several hypotheses have been proposed for thiazide-induced glucose intolerance, including decreased peripheral (46-48) or hepatic insulin sensitivity (49, 50), activation of the sympathetic nervous and the renin angiotensin system by thiazides (11) or attenuation of insulin secretion due to thiazide-induced hypokalemia (51). In our acute model, insulin sensitivity was not affected by HCT and not different between WT and CA5b KO mice. Also, the effect of thiazides on insulin secretion was independent of K^+^. In support of this, a well-designed RCT revealed no significant correlation between changes in plasma K^+^ and serum glucose or plasma insulin in patients treated with HCT, although a significant increase of serum glucose was noted in participants treated with HCT (8). In addition, the profound hypokalemia observed in patients with hyperaldosteronism is not associated with hyperglycemia (52, 53). We did not assess the impact of chronic thiazide administration on glucose metabolism. It is possible that additional mechanisms, including decreased insulin sensitivity or hypokalemia, contribute to thiazide-induced glucose intolerance during long-term administration of thiazides. Clearly, however, acute thiazide administration results in CA5b-dependent attenuation of insulin secretion with subsequent glucose intolerance. Furthermore, chronic mitochondrial CA5b deficiency in β-cells cannot be fully compensated. While the basal insulin secretion is increased, islets of mice with a constitutive CA5b deletion secrete significantly less insulin when stimulated with glucose or sulfonylureas compared to islets of WT mice.

In summary, our results demonstrate that thiazides induce glucose intolerance by an attenuation of insulin secretion in β-cells through inhibition of mitochondrial CA5b.

## METHODS

### Animal studies

Mice were housed 5 per cage in a 12-hour light-dark cycle at 22°C, allowed free access to water, and fed regular rodent chow (#3436 from Provimi Kliba AG, Kaiseraugst, Switzerland).

NCC KO mice (B6.Slc12a3^tm1Ges^), generated by Schultheis et al. (54), were obtained from J. Loffing, University of Zürich, Switzerland and were backcrossed > 10 generations into C57BL/6J background (55). NDCBE KO mice (B6;129S1.Slc4a8^tm1.1Doel^), generated by Leviel et al. (17), were obtained from D. Eladari, Université Pierre et Marie Curie, Paris and were on a mixed C57BL/6J / SV129 background. CA5b KO mice (B6;129S1.Car5b^tm1Sly^), generated by Shah et al. (40), were obtained from Jackson Laboratories (stock # 25570) and were on a mixed C57BL/6J / SV129 background.

For urine collections, mice were housed in individual metabolic cages (Techniplast, Italy). Prior to collections, mice were allowed 2 days to adapt to the metabolic cages. HCT (Sigma-Aldrich, St. Louis, MO) or vehicle (1:1 mixture of 0.9% NaCl solution with DMSO) was applied as a single intraperitoneal injection and urine was collected for the next 6 h (56).

### Intraperitoneal glucose (IPGTT) and insulin (IPITT) tolerance tests

Tolerance tests were performed in 3 months old male mice after a 6 to 12 AM 6 h fast (IPGTT) or at random fed state at 2 PM (IPITT), as described (57-59). Blood glucose was measured at time – 30, 0, 15, 30, 60 and 120 min with a Contour glucose monitor (Bayer Healthcare, Germany) by tail vein sampling in duplicates. Vehicle or HCT was applied by intraperitoneal injection at time -30. Glucose (IPGTT; 2 g/kg, Sigma-Aldrich) or insulin (IPITT; 0.5 or 1 U/kg Actrapid HM, Novo Nordisk, Denmark) were applied by intraperitoneal injection at time 0. For serum insulin measurements tail vein blood sampling was performed at time -30, 0 and 2 min. Vehicle or HCT was applied at time -30 min and glucose (2 g/kg) was applied at time 0 min. Serum insulin was measured with the ultra-sensitive mouse insulin ELISA (CrystalChem, Downers Grove, IL; # 90080).

### Isolation of islets and *in vitro* insulin secretion assays

Pancreata were perfused *in situ* with collagenase solution and islets isolated exactly as described (59, 60). After overnight incubation in RPMI medium with 11 mM glucose, islets were washed twice with KRBH containing (in mM): 115 NaCl, 5 KCl, 25 NaHCO_3_, 0.5 NaH_2_PO_4_, 1.2 MgSO_4_, 2.5 CaCl_2_, 2 glucose, 10 HEPES pH 7.4 and 0.1 % (w:v) BSA. In case of experiments without CO_2_/HCO_3_^-^, the incubation buffer contained (in mM): 140 NaCl, 5 KCl, 0 NaHCO3, 0.5 NaH2PO4, 1.2 MgSO_4_, 2.5 CaCl2, 2 glucose, 10 HEPES pH 7.4 and 0.1 % (w:v) BSA. After washing, islets were placed in 12-well plates (10 islets/well) containing 1 ml of buffer with 2 mM glucose and pre-incubated for 2.5 hrs at 37 °C. Insulin secretion into the supernatant was then measured for 2 h for all secretagogues except for KCl, which was for 1 h. Supernatants were then harvested, plates put on ice and total cellular insulin extracted by addition of acid ethanol (70 % ETOH, 1.5% HCl conc). Secreted and cellular insulin were determined with the ultra-sensitive mouse insulin ELISA.

### Min6 cells

MIN6B1 cells (61), herein referred to as Min6 cells, were provided by Dr. Philippe Halban (University of Geneva, Switzerland) with permission from Dr. Jun-ichi Miyazaki (University of Osaka, Japan) who produced the maternal Min6 cell line (62) and grown in high glucose DMEM supplemented with 10 % heat-inactivated FBS and 70 μM β-mercaptoethanol. Upon receipt and repeatedly thereafter, Min6 cells were tested for mycoplasma contamination. Transfection of Min6 cells with siRNA was done using Lipofectamine 2000 according to the manufacturer’s instruction by a reverse transfection protocol with 0.5 × 10^6^ cells /12-well plate (Invitrogen, Carlsbad, CA). SiRNAs targeting murine CA isoforms were obtained from Quiagen, Hilden. Germany. The following siRNAs were used: CA2 (SI02735992), CA3 (SI02731799), CA5b (SI00941157), CA6 (SI00941192), CA7 (SI02745442), CA8 (SI00167601), CA9 (SI00235340), CA10 (SI00941094), CA11 (SI00187754), CA13 (SI00222649), CA15 (SI02719780).

72 h after transfection, cells were washed 3x with Krebs-Ringer bicarbonate HEPES buffer (KRBH) containing (in mM): 135 NaCl, 3.6 KCl, 5 NaHCO_3_, 0.5 NaH_2_PO_4_, 0.5 MgCl_2_, 1.5 CaCl_2_, 2 glucose, 10 HEPES pH 7.4 and 0.1 % (w:v) BSA. In case of experiments without CO_2_/HCO_3_^-^, the incubation buffer contained (in mM): 140 NaCl, 5 KCl, 0 NaHCO3, 0.5 NaH2PO4, 1.2 MgSO_4_, 2.5 CaCl2, 2 glucose, 10 HEPES pH 7.4 and 0.1 % (w:v) BSA. After an equilibration period of 2 h at 37 °C, cells were incubated for 1 h in fresh KRBH buffer containing either 2 mM glucose or indicated secretagogues, for KCl stimulation incubation time was 15 min (61). After this period, supernatants were harvested, plates put on ice and total cellular insulin extracted by addition of acid ethanol (70 % ETOH, 1.5% HCl conc). Secreted and cellular insulin were determined with the ultra sensitive mouse insulin ELISA. Oxalacetate was quantified by a colorimetric assay (Abcam, Cambridge, UK; # ab83428).

### Cell viability assay

Cell viability was assessed using the colorimetric MTS assay (Promega, Madison, WI; # G8080) according to the manufacturer’s instructions. Min6 cells or islets were treated as described for glucose-stimulated insulin secretion experiments in KRBH buffer, and incubated with the vehicle DMSO (1:1000) or indicated thiazide diuretics for 4 h prior to the assessment of cell viability.

### Measurement of pyruvate carboxylase activity

Pyruvate carboxylase was measured in Min6 or islet lysates using the colorimetric pyruvate carboxylase activity assay kit (Biovision, CA, USA; # K2075-10) according to the manufacturer’s instructions. Prior to cell lysis, cells were cultured in CO_2_/HCO_3_^-^ free conditions for 4 h.

### RNA Isolation, RT-PCR, and quantitative real-time PCR

Total RNA was isolated using Trizol® reagent as detailed in the manufacturer’s protocol (Invitrogen, Carlsbad, CA). Reverse transcription was performed using the Taqman® Reverse Transcription kit (Life Technologies/ABI, Rotkreuz, CH). Real-time PCR was performed using presynthesized Taqman®-based Assays-on-Demand (Life Technologies/ABI) on an ABI ViiA 7 System. The following Assays-on-Demand were employed: Slc12a3 (NCC) (Mm00490213_m1), SLC4A8 (NDCBE) (Mm00491397_m1), SLC12A2 (NKCC1) (Mm01265951_m1), SLC12A1 (NKCC2) (Mm01275821_m1), SLC26A4 (Pendrin) (Mm01258316_m1), CA1 (Mm00486717_m1), CA2 (Mm00501576_m1),CA3 (Mm01281795_m1), CA4 (Mm00483021_m1), CA5a (Mm00500371_m1), CA5b (Mm00490359_m1), CA6 (Mm00486722_m1), CA7 (Mm01247656_m1), CA8 (Mm00801469_m1), CA9 (Mm00519870_m1), CA10 (Mm01304922_m1), CA11 (Mm00515393_m1), CA13 (Mm01291526_m1), CA14 (Mm00488564_m1), CA15 (Mm00472660_m1), GAPDH (Mm9999915_g1). Ct values for triplicate technical replicates were averaged, and the amount of mRNA relative to GAPDH was calculated using the ΔCt method.

### Statistical analysis

Data are presented as mean ± SD. For comparisons between groups the 2-tailed Student’s t test (two groups) or 1-way ANOVA with Tukey post-hoc test (multiple groups) was used.

Data analysis was done with GraphPad Prism 9.3.1 (GraphPad Software, San Diego, CA, USA). All statistical tests were two-sided and p < 0.05 was considered statistically significant.

### Study approval

All animal experiments were in accordance with the Swiss Animal Welfare Law and approved by the Local Veterinary Authority of the Kanton Bern (permits # 21/14, 116/17, 110/19) and complied with the ARRIVE guidelines.

## Author contributions

DGF and DK conceptualized the study and acquired financial support. PK, SW, DK and DGF designed experiments. PK, GA, CD performed experiments. PK, SW, DK and DGF performed data analysis and data interpretation. PK and DGF wrote the first draft of the manuscript. All authors contributed to discussion and editing of text and approved the final version of the manuscript.

## Data availability

All raw data, processed data and corresponding metadata will be made available on Dryad (https://datadryad.org/stash) upon publication of the manuscript.

## Acknowledgements

DGF was supported by the Swiss National Science Foundation (# grant 31003A_172974), the Swiss National Centres of Competence in Research (NCCR TransCure and NCCR Kidney.CH), the Novartis Research Foundation and by a Medical Research Position Award of the Foundation Prof. Dr. Max Cloëtta. We thank D. Eladari and J. Loffing (transgenic mice) and P. Halban (Min6 cells).

