## Supplemental Information for "Thiazides induce glucose intolerance through inhibition of mitochondrial carbonic anhydrase 5b in β-cells"

**SUPPLEMENTAL FIGURES**

**Supplemental Figure 1**

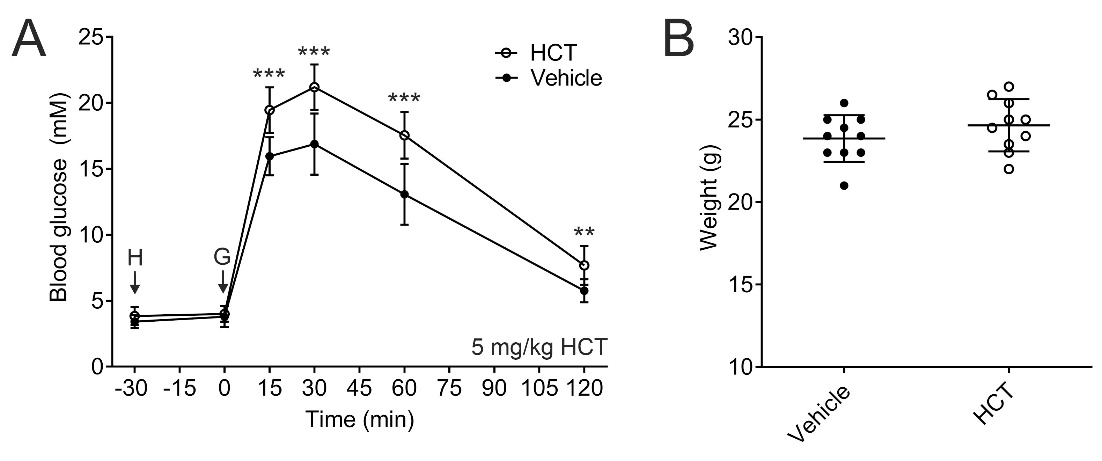

**Glucose tolerance test with hydrochlorothiazide** **treatment after overnight fasting.** Intraperitoneal glucose tolerance tests **(A)** and corresponding body weights **(B)** of C57BL/6J mice treated with 5 mg/kg hydrochlorothiazide (HCT) or vehicle after 16 h overnight fasting. HCT or vehicle i.p. (arrow H) was applied at time point -30 min, glucose (2g/kg; arrow G) was applied at time point 0 min. Whole blood glucose was measured at indicated time points. N=10 mice per group. Data are show as mean ± SD. Asterisks denote significance for comparisons between groups of mice at indicated time points (two-tailed unpaired Student's t-test; **p < 0.01, ***p < 0.001).

**Supplemental Figure 2**

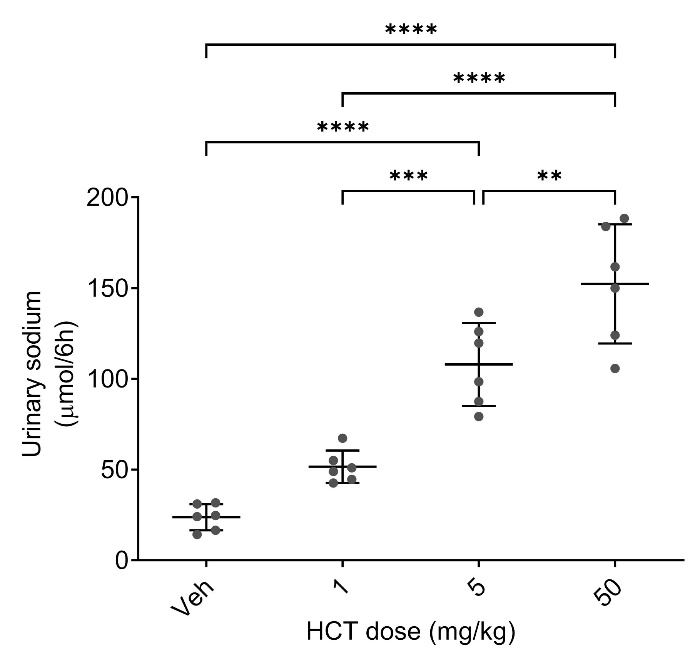

**Dose-dependence of the natriuretic effect of hydrochlorothiazide in mice.** Individually in metabolic cages housed C57BL/6J mice received indicated dose (mg/kg body weight) of hydrochlorothiazide (HCT) or vehicle (Veh) i.p. Depicted is the cumulative urinary sodium excretion in the 6 h after application of HCT or vehicle. Each dot represents the measurement in an individual mouse, data are show as mean ± SD. N=6 mice per treatment group. Asterisks denote significance for the indicated comparisons (ANOVA with Tukey post-hoc test; **p < 0.01, ***p < 0.001, ****p < 0.0001).

**Supplemental Figure 3**

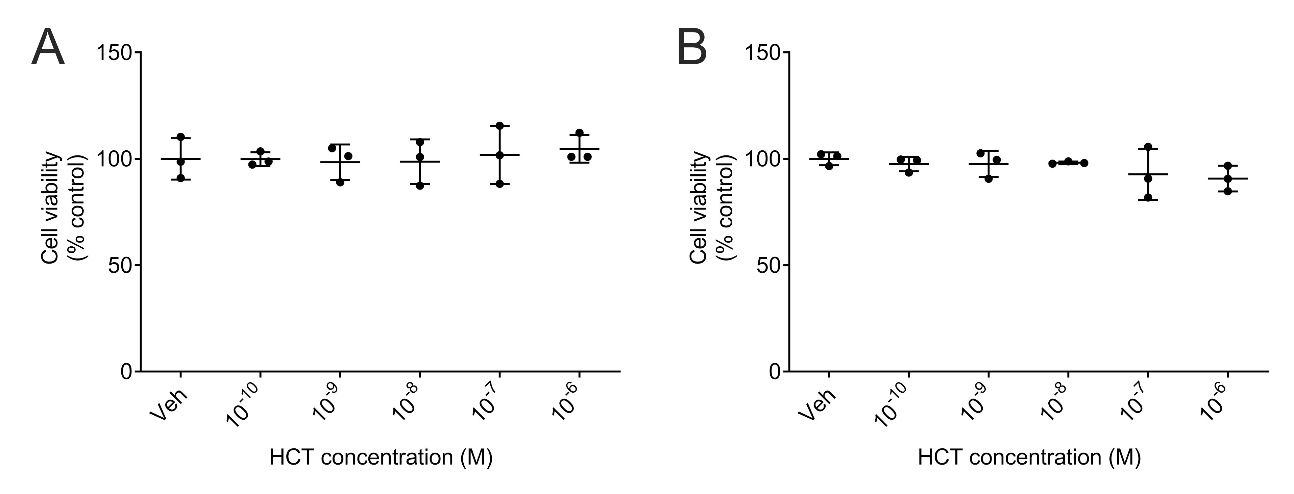

**Assessment of cell viability in murine islets or Min6 cells treated with hydrochlorothiazide.** Cell viability in murine islets **(A**) or Min6 cells **(B)** assessed by the MTS assay in the presence of indicated concentrations of hydrochlorothiazide (HCT) normalized to the vehicle (Veh) treated condition. Islets and Min6 cells were exposed to HCT or Veh for 4 hrs . Each dot represents islets isolated of an individual mouse. In case of Min6 cells, each dot represents an individual experiment Data are shown as individual observations with mean ± SD. No significant differences between groups were detected (ANOVA with Tukey post-hoc test).

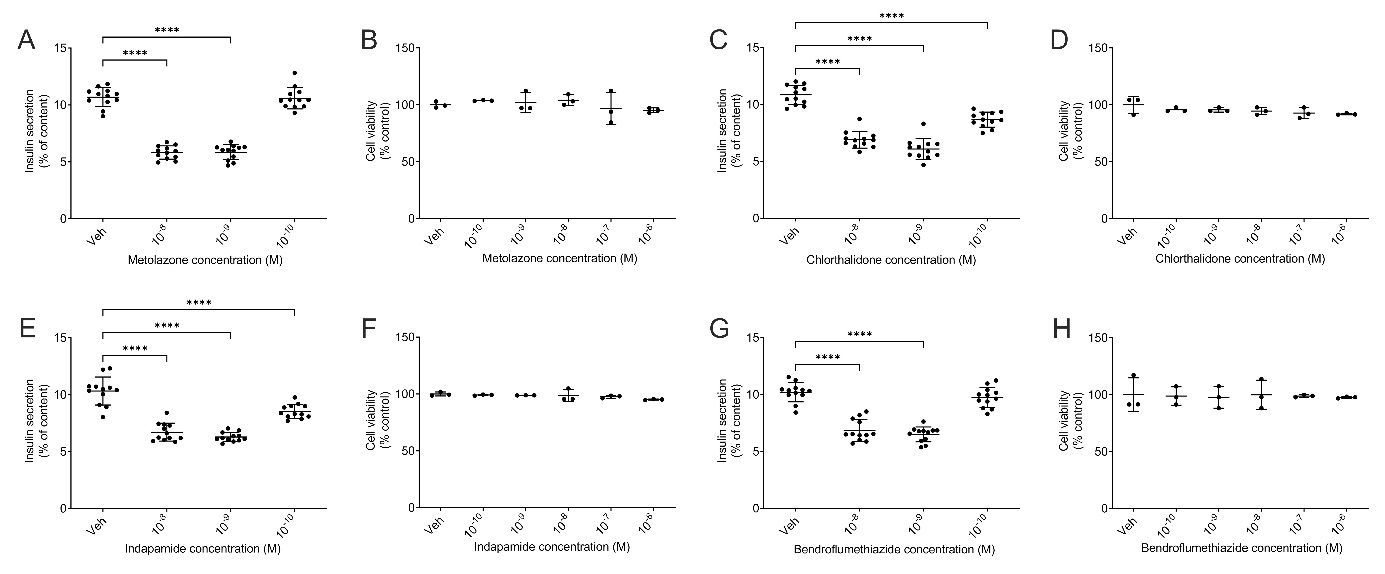
**Supplemental Figure 4**

**Effect of thiazide diuretics and thiazide-like diuretics on insulin secretion.** Glucose-stimulated insulin secretion (20 mM glucose; GSIS) and cell viability (MTS assay) in Min6 cells incubated with vehicle (Veh) or metolazone **(A** and **B)**, chlorthalidone **(C** and **D)**, indapamide **(E** and **F)** or bendroflumethiazide **(G** and **H)** at indicated concentrations. Data represent three individual experiments combined. Data are shown as individual observations with mean ± SD. Asterisks denote significance for the indicated comparisons (ANOVA with Tukey post-hoc test; ****p < 0.0001).

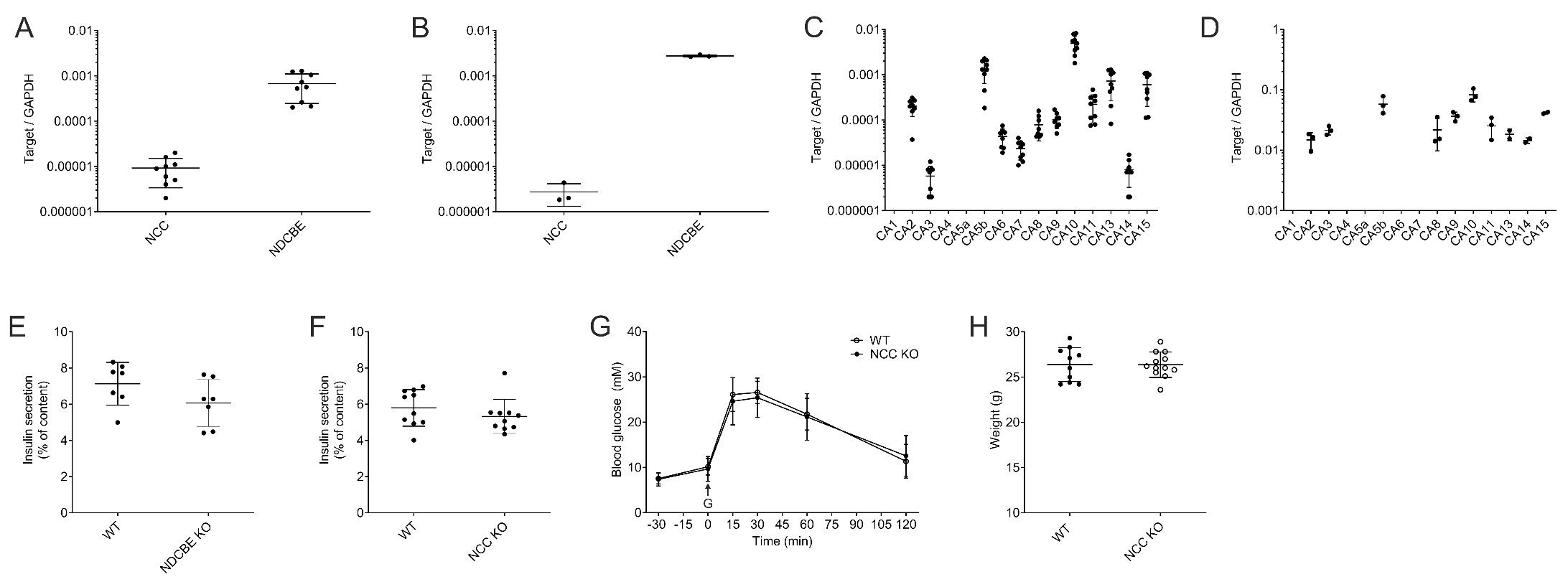
**Supplemental Figure 5**

**Expression of thiazide targets in Min6 cells and islets and insulin secretion studies of islets isolated from NCC and NDCBE KO mice.** Expression of NCC (SLC12A3) and NDCBE (SLC4A8) transcripts in Min6 cells **(A)** and murine islets **(B)** quantified by real-time PCR normalized to GAPDH. Expression of CA isoforms 1 - 15 in Min6 cells **(C)** and murine islets **(D)** quantified by real-time PCR normalized to GAPDH. Min6 cell experiments represent three individual experiments combined **(A** and **C)**. In case of islet expression studies, each dot represents islet mRNA isolated from an individual mouse **(B** and **D)**. Data are shown as individual observations with mean ± SD.

Glucose-stimulated insulin secretion (20 mM glucose; GSIS) of islets isolated from WT and NDCBE **(E)** or WT and NCC **(F)** mice. Each dot represents islets isolated of an individual mouse. **(G** and **H)** Intraperitoneal glucose tolerance tests and corresponding body weights of WT and NCC KO mice. Glucose (2g/kg i.p.; arrow G) was applied at time point 0 min. Whole blood glucose was measured at indicated time points. Comparisons between the two groups of mice at times of measurement **(G)** or comparison of weights of the two groups of mice **(H)** with two-tailed unpaired Student's t-test ; no significant differences were observed. N=10 WT and 12 NCC KO mice.

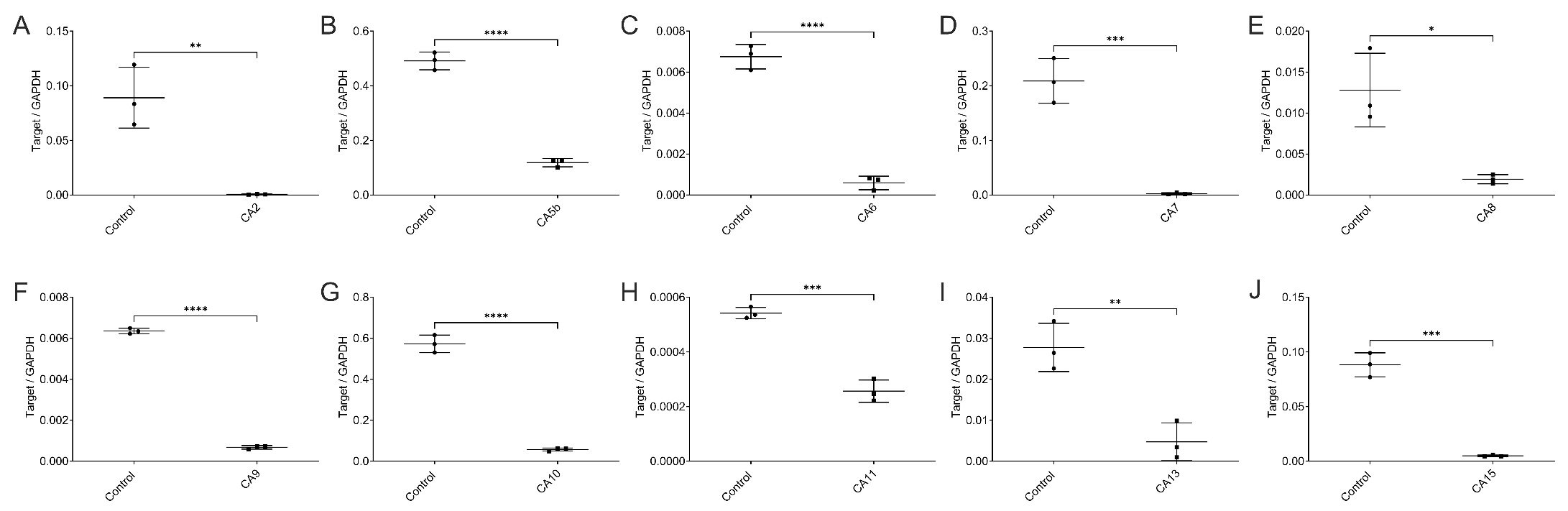
**Supplemental Figure 6**

**Knock-down of CA isoforms in Min6 cells by siRNA.** Min6 cells were treated with control siRNA or siRNA targeting the indicated CA isoform **(A** - **J)** and CA isoform mRNA expression was quantified by real-time PCR and normalized to GAPDH mRNA expression. Results show three individual experiments combined. Data are shown as individual observations with mean ± SD. Asterisks denote significance for the indicated comparisons (ANOVA with Tukey post-hoc test; *p < 0.05, **p < 0.01, ***p < 0.001, ****p < 0.0001).

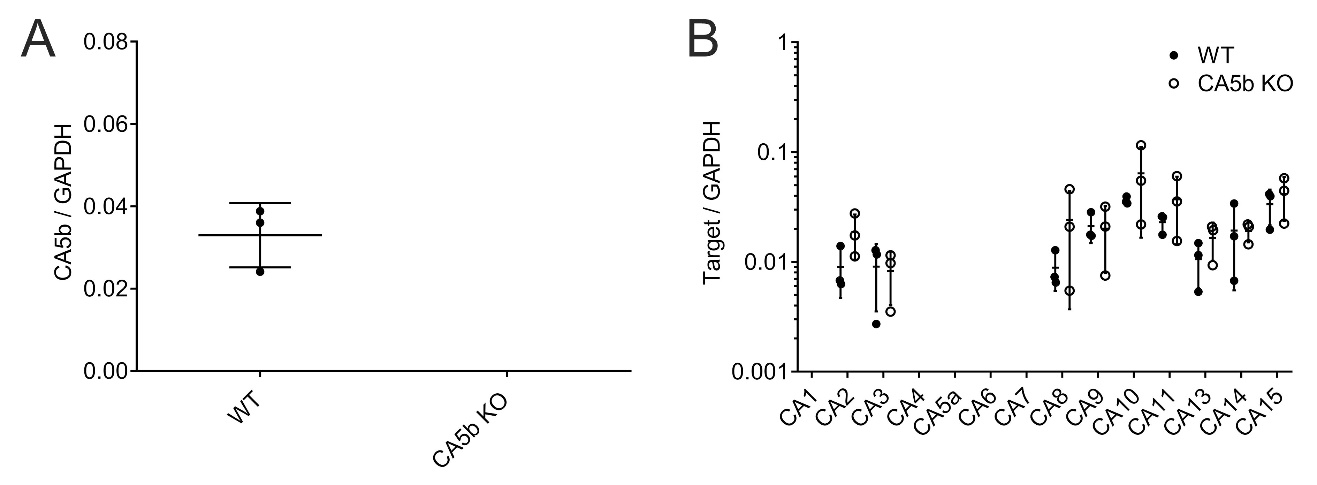
**Supplemental Figure 7**

**CA isoform expression in islets of WT and CA5b KO mice. (A)** Expression of CA5b mRNA in islets isolated of WT and CA5b KO mice quantified by real-time PCR normalized to GAPDH. No CA5b transcript was detectable in CA5b KO mice. (B) Expression of CA isoform mRNA expression in islets isolated of WT and CA5b KO mice quantified by real-time PCR normalized to GAPDH. Each dot represents islet mRNA isolated from an individual mouse. No difference in CA isoform expression was detectable between WT and CA5b KO islets (ANOVA with Tukey post-hoc test). Data are shown as individual observations with mean ± SD.
